# Efficient genotype compression and analysis of large genetic variation datasets

**DOI:** 10.1101/018259

**Authors:** Ryan M. Layer, Neil Kindlon, Konrad J. Karczewski, Exome Aggregation Consortium, Aaron R. Quinlan

**Affiliations:** Departments of Human Genetics and Biomedical Informatics, University of Utah, Salt Lake City, UT; Department of Public Health Sciences, University of Virginia, Charlottesville, VA; Analytical and Translational Genetics Unit, Harvard Medical School, Boston, MA

## Abstract

The economy of human genome sequencing has catalyzed ambitious efforts to interrogate the genomes of large cohorts in search of new insight into the genetic basis of disease. This manuscript introduces Genotype Query Tools (GQT) as a new indexing strategy and toolset that addresses an analytical bottleneck by enabling interactive analyses based on genotypes, phenotypes and sample relationships. Speed improvements are achieved by operating directly on a compressed genotype index without decompression. GQT’s data compression ratios increase favorably with cohort size and relative analysis performance improves in kind. We demonstrate substantial performance improvements over state-of-the-art tools using datasets from the 1000 Genomes Project (46 fold), the Exome Aggregation Consortium (443 fold), and simulated datasets of up to 100,000 genomes (218 fold). Furthermore, we show that this indexing strategy facilitates population and statistical genetics measures such as principal component analysis and burden tests. Based on its computational efficiency and by complementing existing toolsets, GQT provides a flexible framework for current and future analyses of massive genome datasets.

## INTRODUCTION

For the majority of common human diseases, only a small fraction of the heritability can be explained by the genetic variation we know of thus far. One hypothesis posits that the elusive heritability is explained in part by rare, and thus largely unknown, genetic variation in the human population^1^. Multiple efforts are therefore underway to sequence hundreds of thousands of human genomes in order to catalog the full spectrum of human genetic variation from the common to the vanishingly rare. Extrapolating from current and forthcoming efforts, it is likely that more than 1 million human genomes will be sequenced in the near term. Integrated analyses and community sharing of such population datasets will be critical for future discovery. However, in aggregate, the resulting datasets will include roughly 100 trillion distinct genotypes from the more than 100 million polymorphic loci that are likely to be discovered. Therefore, the development of more effective data compression and exploration strategies will be crucial to enable discovery and to make these valuable datasets available to all researchers.

The need for computationally efficient representations of genotype datasets is not new. In 2007, Purcell et al. introduced the binary PED (BED) format for the popular PLINK variant association testing software^2^. The BED format encodes four biallelic genotypes in each byte using two bits per individual genotype. Binary encoding reduces the size of the resulting data file versus a simple text representation. The more recent Variant Call Format (VCF)^3^ represents variants, sample genotypes, and flexible variant annotations from DNA sequencing studies and its binary counterpart (BCF) provides a more efficient means of storing such datasets via compression and highly structured data. The BED and VCF/BCF formats, as well as their associated toolsets have become the standard for genome variation research. However, because these formats are intentionally structured from the perspective of polymorphic loci in order to support “ variant-centric” analyses (e.g., which variants overlap BRCA1?), they are consequently ill suited to queries driven by specific genotype and phenotype combinations, inheritance patterns, or allele frequency thresholds (e.g., which variants are enriched in affected daughters?). Since detailed knowledge of cohort ancestry, phenotype and family structure are fundamental to interpreting genetic architecture of disease, the need for new analysis methods is increasingly acute as cohort sizes increase.

Here we introduce a new, “ individual-centric” strategy for indexing and mining individual genotypes and variants ascertained from millions of genomes. Our approach, which is made freely available in the Genotype Query Tools (GQT) software package, reorganizes and indexes genotype data such that it optimizes queries screening for variants based on the genotypes, allele frequencies, and phenotypes of one or more of the individuals in the study. GQT also utilizes an efficient data compression strategy to minimize the disk storage requirements of its index. Our goal is to provide a genotype compression and indexing scheme that maximizes analysis performance by allowing direct measurement and comparison of the compressed data without inflation. We demonstrate that these improvements provide extremely compact genotype compression and efficient queries based on sample genotypes and phenotypes that are orders of magnitude faster than existing methods. We further illustrate how the GQT strategy can be used to explore large-scale datasets and quickly compute common population and statistical genetics measures.

## RESULTS

### Overview of the GQT genotype indexing strategy

Because existing formats such as VCF (Figure 1A) are organized to support variant-centric analyses, individual-centric data exploration based on sample genotypes, phenotypes and relationships is inherently inefficient, since every variant row must be tested in search of variants meeting specific genotype criteria (Figure 1B). GQT overcomes these limitations by creating a compressed, individual-centric index, as well as a powerful query interface to improve both speed and analytical flexibility. Speed improvements are achieved by operating directly on the compressed index without inflation, and query flexibility is driven by combining sample metadata (phenotypes, population, gender, etc.) with genotype filters in an integrated query interface.

**Figure 1.**
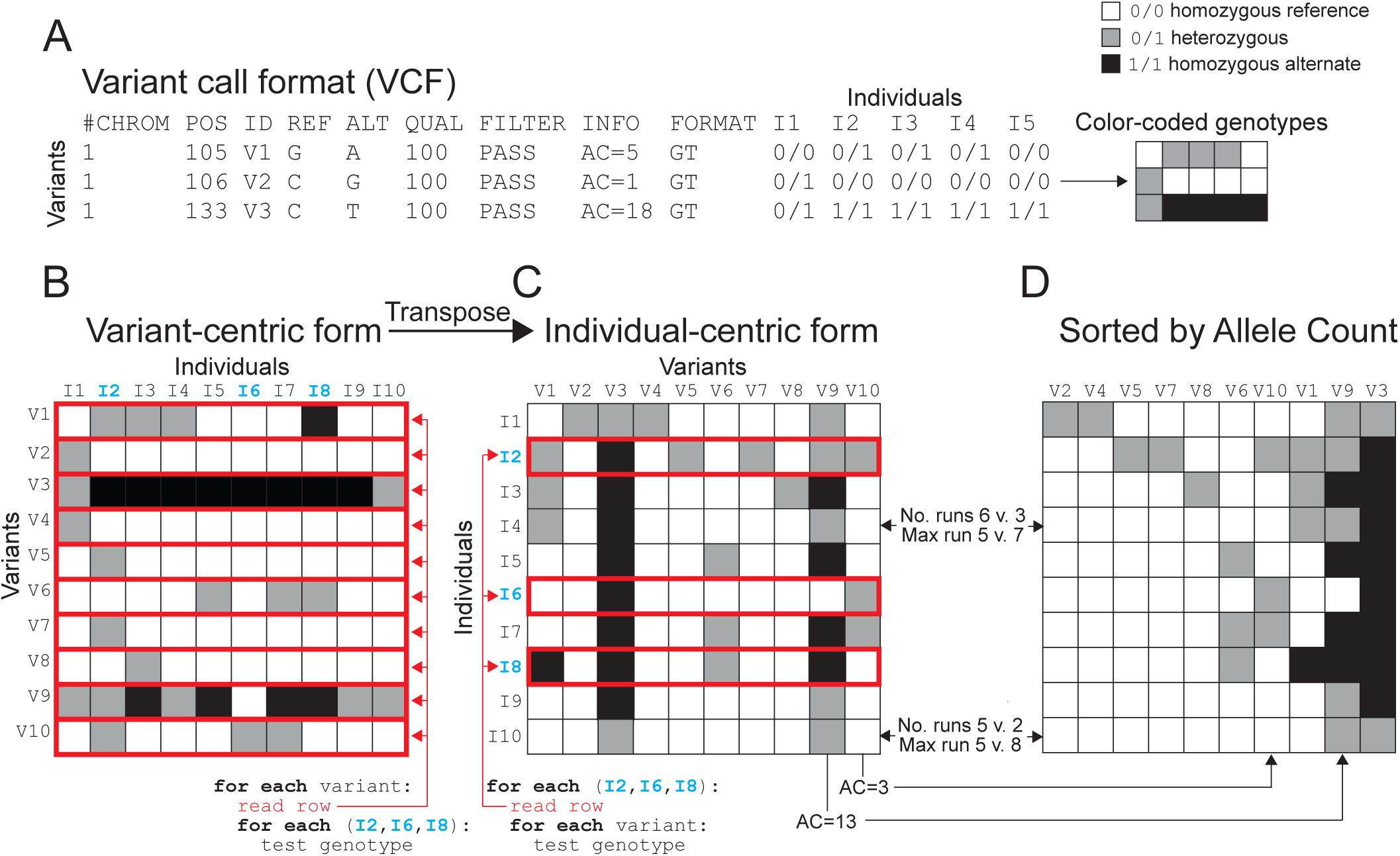
Individual-centric data organization. (**A**) The “ variant-centric” VCF standard is essentially a genotype matrix where rows correspond to variants and columns to individuals. (**B**) The VCF standard is inefficient for queries across all genotypes and a subset of individuals since each variant row must be inspected to test all of the genotypes of specific individuals. (**C**) By transposing the matrix such that rows (data records) now represent the full set of genotypes for each individual, the data better aligns to individual-centric questions and algorithms. (**D**) Sorting the columns of an individual-centric matrix by alternate allele count (AC) improves compressibility.

The GQT genotype index is created by transposing genotypes from the VCF format’s variant-centric form (Figure 1B; each row is a variant) to an individual-centric form (Figure 1C; each row is now an individual), which minimizes the number of rows that must be accessed when comparing the genotypes of the individuals in a dataset. Variant columns are then sorted by allele frequency (Figure 1D) to take advantage of the fact that the majority of genetic variation is extremely rare in the population^4,5^. Reordering by allele frequency greatly improves data compression since it yields much longer runs of identical genotypes than transposition alone (Supplementary Figure 1). After sorting by allele frequency, each individual’s genotypes are encoded into a bitmap index comprised of four distinct bit arrays corresponding to each of the four possible (including “ unknown”) diploid genotypes (Supplementary Figure 2). Bitmaps allow efficient comparisons of many genotypes in a single operation using bitwise logical operations, thereby enabling rapid comparisons of sample genotypes among many variants in the original VCF file (Supplementary Figure 3). Lastly, the bitmap indices are compressed using Word Aligned Hybrid (WAH) encoding^6^, which achieves near-optimal compression while still allowing bitwise operations directly on the compressed data (see Supplementary Figure 4). This encoding minimizes the disk storage requirements of the index and, by eliminating the need for data inflation, improves query speed since runtime is a function of the compressed input size.

### GQT index scalability

GQT’s genotype index is designed for rapid individual-centric query performance, and is intended to be a complement to the original VCF file and its variant-centric index. Therefore, the GQT index requires some additional (albeit minor) storage cost. Additionally, returning the variants that match a query from the VCF file quickly becomes a bottleneck since existing VCF toolkits cannot return arbitrary (e.g., the million) variant records from a VCF file. GQT therefore creates two additional indices (BIM and VID) that allow variants satisfying an analysis to be quickly returned in VCF format (see Supplementary Figure 5). The storage cost of the GQT indices is marginal with respect to the size of the underlying VCF file and its size continues to diminish as the cohort size grows since most new variation discovered will be very rare. To demonstrate this, we compare the GQT index and PLINK BED encoding of the Phase 3 of the 1000 Genomes project (2,504 individuals and 84,739,846 variants) to the size of the uncompressed VCF for the full dataset (Figure 2A). We also include the size of a BCF compressed version of the 1000 Genomes VCF as a point of comparison for the size of the GQT index.

**Figure 2.**
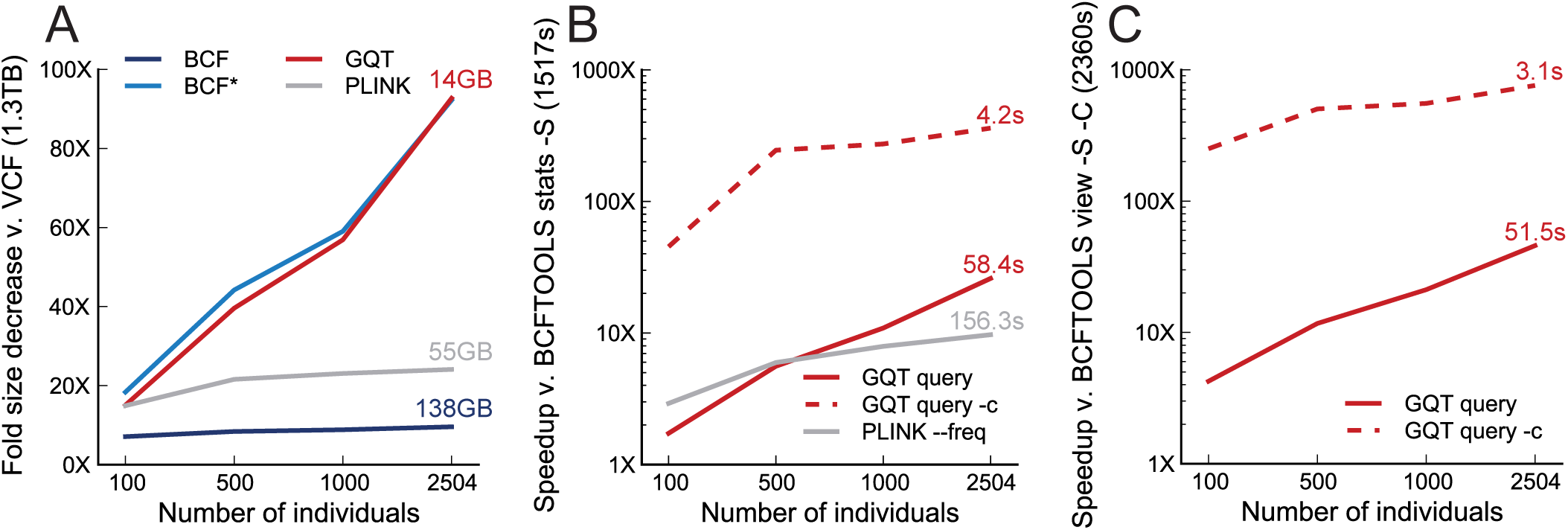
GQT compression and query performance for 1000 Genomes data (Phase 3). (**A**) File size reduction for 1000 genomes phase 3, which is comprised of 2,504 individuals and over 84 million variants. Compression ratios describe the fold reduction in file size relative to an uncompressed VCF file. GQT reflects the total size of the GQT genotype index plus the size of the BIM and VID indices (see Supplement for details). BCF represents the size of the full BCF file including genotype metadata, whereas BCF* omits genotype metadata. Lastly, PLINK reflects the size of PLINK v1.9 BED and BIM files. (**B**) Fold speedup for computing the alternate allele frequency count for a targeted subset of 10% of the 2,504 individuals. The baseline for comparison was the BCFTOOLS “`stats`” command. Two versions of GQT output are considered, all matching variants in VCF (GQT query), and the matching variants (GQT query –c). (**C**) Fold speedup for finding variants having an allele frequency of less than 1% in a target 10% of individuals. The baseline for comparison was BCFTOOLS “`view -C".` Note that PLINK v1.9 was excluded from this comparison because it does not directly compute this operation.

Both the BCF and PLINK encodings exhibited constant compression across population sizes with a 9.6X improvement (138.4 GB) and 24.1X reduction (55.1 GB), respectively, for the full 2,504 individuals. In contrast, the reduction in the relative size of the GQT indices steadily improved as the number of individuals and variants increased, yielding a 92.7X (14.3 GB) reduction for 2,504 individuals and requiring, on average, only 0.54 bits per genotype. The BCF encoding of the 1000 Genomes dataset included extensive metadata such as genotype likelihoods and allelic read depths for each individual and variant. It is encouraging that when we removed this metadata (BCF* in Figure 2A) in order to provide a direct comparison to the storage requirements for all of the GQT indices, the compression rates were nearly identical. This demonstrates that for genotypes from a large human cohorts, the GQT compression strategy is on par with the LZ77 algorithm and it is not sacrificing additional compression.

### GQT query interface

We have created software that provides analytical power by leveraging the GQT index to address a broad range of questions and analyses. This toolset provides a simple interface for variant queries based on sample genotypes and metadata (Figure 3). GQT creates an SQLite database from a PED file describing the names, relationships, multiple phenotypes, and other attributes of the samples in an associated VCF file (Figure 3A). The sample database complements the GQT index of the original VCF/BCF file and allows GQT to identify the rows of sample genotypes in the GQT index that match the query. For example, Figure 3B demonstrates a hypothetical GQT query in search of variants where all affected individuals (“`phenotype==2`”) are heterozygous. Using these criteria, the GQT query tool issues the appropriate SQL query to the sample database to retrieve the position of genotype bitmaps of three affected individuals within the GQT index. Once identified, the bit arrays for heterozygous genotypes are AND’ed to isolate the variants where all three individuals are heterozygous.

**Figure 3.**
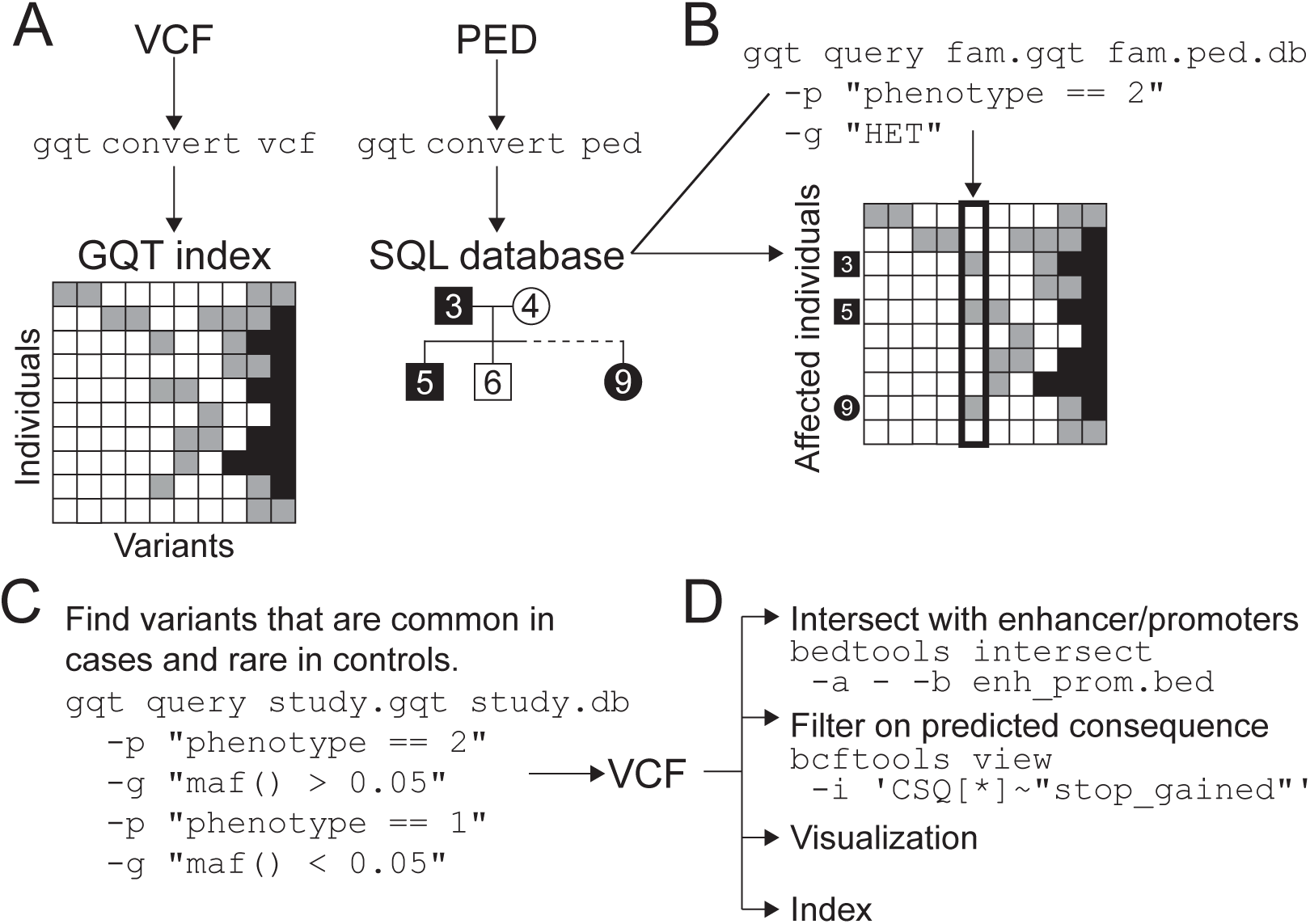
GQT enables queries based on sample genotypes and phenotypes. **(A)** GQT creates an index of the sample genotypes in a VCF or BCF file. In addition, GQT will create a SQLite database of a PED file describing the familial relationships, gender, ancestry, and various, user-defined phenotypes of the samples in the VCF file. (**B**) The PED database allows GQT to quickly find the specific sample genotype rows in the genotype index that correspond to the samples that meet a user’s search criteria. For example, a search for variants where all affected individuals are heterozygous begins with a query to the PED database, which identifies samples 3,5 and 9 as affected. In turn, the GQT index is used to quickly find the sole variant where samples 3,5, and 9 are all heterozygous. (**C**) Providing multiple query conditions can further refine variant searches. (**D**) Since GQT returns the identified rows in VCF format, it can be combined with other, variant-centric tools such as BEDTOOLS and BCFTOOLS to enable sophisticated analyses based on sample genotypes, variant properties, and genome annotations. This also supports visualization or variant-centric indexing with tools such as TABIX.

Queries may be based on any attribute that is defined in the PED file and queries may combine multiple criteria. As examples, one may search for variants having markedly different minor allele frequencies in different world sub-populations or case versus control samples (Figure 3D). Moreover, query results are returned in VCF format, thereby allowing sophisticated analyses combining GQT queries with other, variant-centric tools such as BEDTOOLS^7^, VCFTOOLS^3^ and BCFTOOLS (unpublished).

### Query performance

The typical tradeoff for high data compression is the time spent re-inflating compressed data prior to analysis. We designed our indexing strategy precisely to avoid this tradeoff and achieve efficient queries of cohorts involving thousands to millions of individuals. To demonstrate this, we compared the query performance of GQT to BCFTOOLS and PLINK. First, we compared the time required to compute the alternate allele frequency among a target set of 10% of individuals from the 1000 Genomes VCF (Figure 2B). Whereas BCFTOOLS required 1517.5 seconds, both GQT and PLINK were substantially faster, requiring 58.4 seconds (26.0 fold speedup) and 156.3 seconds (9.6 fold speedup). Importantly, GQT’s performance improved as the number of individuals increased, whereas PLINK’s performance was relatively flat. Moreover, matching variants are identified almost instantly and thus, the majority of GQT’s runtime is spent emitting the VCF results of the query. For example, when the GQT “count” option is invoked to simply return the count of variants matching the query (i.e., without returning the full variant records themselves), the runtime dropped to 4.2 seconds. We also compared the time required to identify rare (AAF<1%) variants among a subset of 10% of the individuals (Figure 2C). In this case, GQT was up to 45.8X faster than BCFTOOLS (51.5 seconds v. 2360.5 seconds). PLINK was not included in the rare variant search comparison because it does not natively support that function.

GQT’s query performance relative to existing methods continues to improve for even larger datasets. When considering the Exome Aggregation Consortium (ExAC) variant dataset (9.36 million exonic among 60,706 human exomes), the GQT index was only 0.2% the size of the VCF (28GB v. 14.1 TB), reflecting a storage requirement of merely 0.38 bits per genotype. Moreover, rare variants were found in only 2.1 minutes (9.98 seconds when excluding the time required to report the variants), reflecting a 443.5 fold improvement over BCFTOOLS (931.4 minutes). Furthermore, based on simulated datasets involving 100 to 100,000 genomes (Methods), it is clear that GQT’s data compression and query performance continues to improve dramatically with larger cohorts. While simulating variants from one million or more individuals is computationally intractable for this study, extrapolation suggests that GQT indices involving a cohort of a million genomes will yield query performance that is at least 218 fold faster than BCFTOOLS (Figure 4).

**Figure 4.**
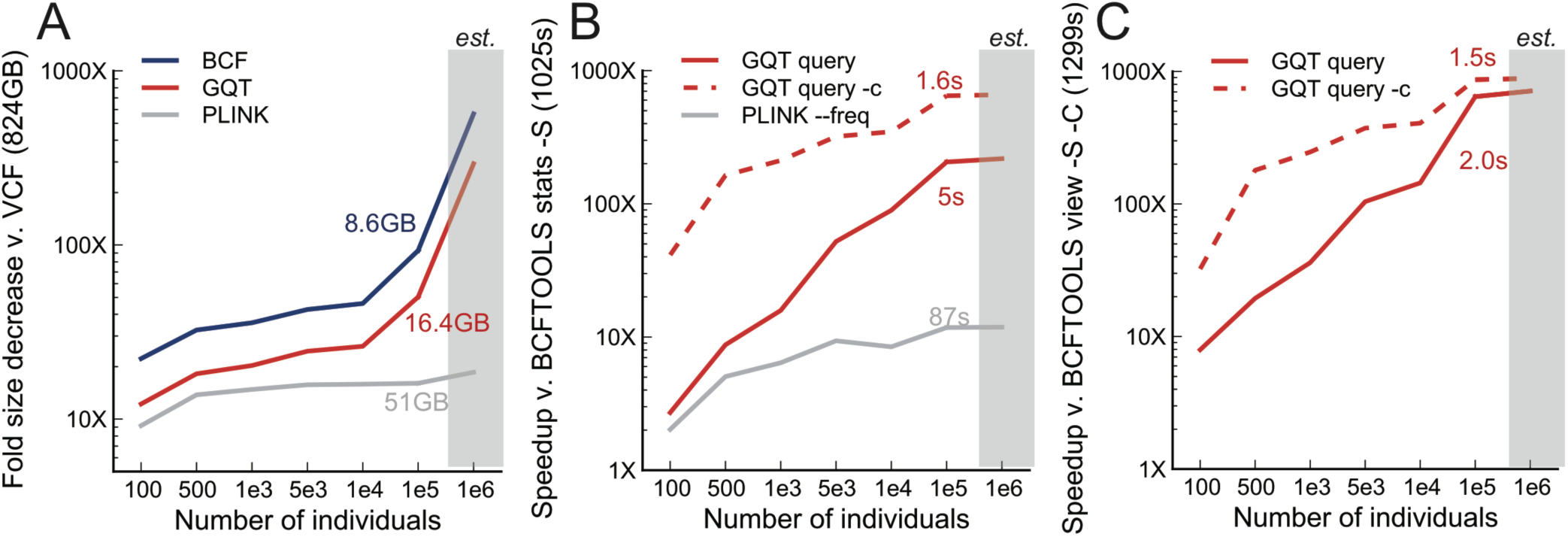
GQT compression and query performance for simulated genomes. A comparison of BCFTOOLS, GQT, and PLINK for simulated genotypes on a 100 Mb genome with between 100 and 100,000 individuals. (**A**) The fold reduction each tool provided relative to the original, uncompressed VCF file. (**B**) Fold speedup for computing the alternate allele frequency count for a target 10% of individuals. (**C**) Fold speedup for finding variants having an allele frequency of less than 1% in a target 10% of individuals. Note that this simulation did not include either variant or sample metadata and that the metrics for 1 million individuals (*est.*) were estimated using a simple linear fit. As in Figure 2C, PLINK was excluded from this comparison because it does not directly compute this operation.

### Applications of the GQT genotype indexing strategy

GQT’s combination of an individual-centric genotype reorganization with a compressed bitmap indexing strategy enables a flexible and efficient query interface for analyses based on sample genotypes and phenotypes. Since this indexing strategy is fundamentally optimized for questions that involve comparisons of sample genotypes among many variant loci, it is also as well suited to expedite many common statistical and population genetics measures. Therefore, we anticipate that GQT genotype indices will empower other statistical genetics software and will serve as a framework for future method development. In the following sections, we present four applications of the GQT indexing strategy to demonstrate its generality and flexibility for a broad range of analyses.

#### Pedigree analyses

Using the GQT query interface, we screened for high confidence de novo mutations in the CEPH 1473 pedigree sequenced as part of the Illumina Platinum Genomes project (Figure 5A; Supplementary Figure 6A). A preliminary search for candidate de novo mutations in the extensively studied NA12878 daughter involves screening for variants that are homozygous for the reference allele in NA12878’s parents, yet are heterozygous in NA12878. GQT identifies 11,172 such candidates from more than 8 million total variants in 0.04 seconds (Supplementary Figure 6B). A more sophisticated GQT query recognizes that true de novo mutations in the germline of NA12878 should be inherited by her offspring, reducing the set of candidates to 3,002 (Supplementary Figure 6C). Excluding suspicious variants that lie in low-complexity regions^8^ reduces the set of de novo mutation candidates by another 12% (N=2,659). While this yields many more candidates than would be predicted by the 1.2 × 10^-8^ per generation base pair mutation rate observed in the CEU pedigree^5^, the 1000 Genomes Project employed additional filters based on genotype likelihoods, proximity other variants, and other properties of the sequence alignments^5^. Moreover, the intent of this analysis is to demonstrate GQT’s analytical power in the context of both large studies of unrelated individuals as well as family-based studies of disease.

#### Principal component analysis (PCA)

PCA is commonly used in genetics to identify population substructure by comparing the genotypes of large cohorts in search of variants that distinguish (i.e., are enriched in) individuals with a particular ancestry. The fundamental complexity with most existing PCA algorithms is that the genotypes of each individual are compared to every other individual’s genotypes to produce pairwise measures of genotype similarity. As such, half a million pairwise genotype comparisons are required to produce a matrix describing the genetic similarity of a cohort of 1000 individuals. Once constructed, common statistical packages are available to identify the principal components from the similarity matrix. Since GQT is optimized for rapid genotype comparisons among individuals, it is especially well suited for PCA. In fact, we are able to compute the similarity matrix for the 2,504 individuals in the Phase 3 1000 Genomes dataset in 207 minutes, and an analysis of solely the 347 admixed American (AMR) individuals required merely 3 minutes (Figure 5B).

**Figure 5.**
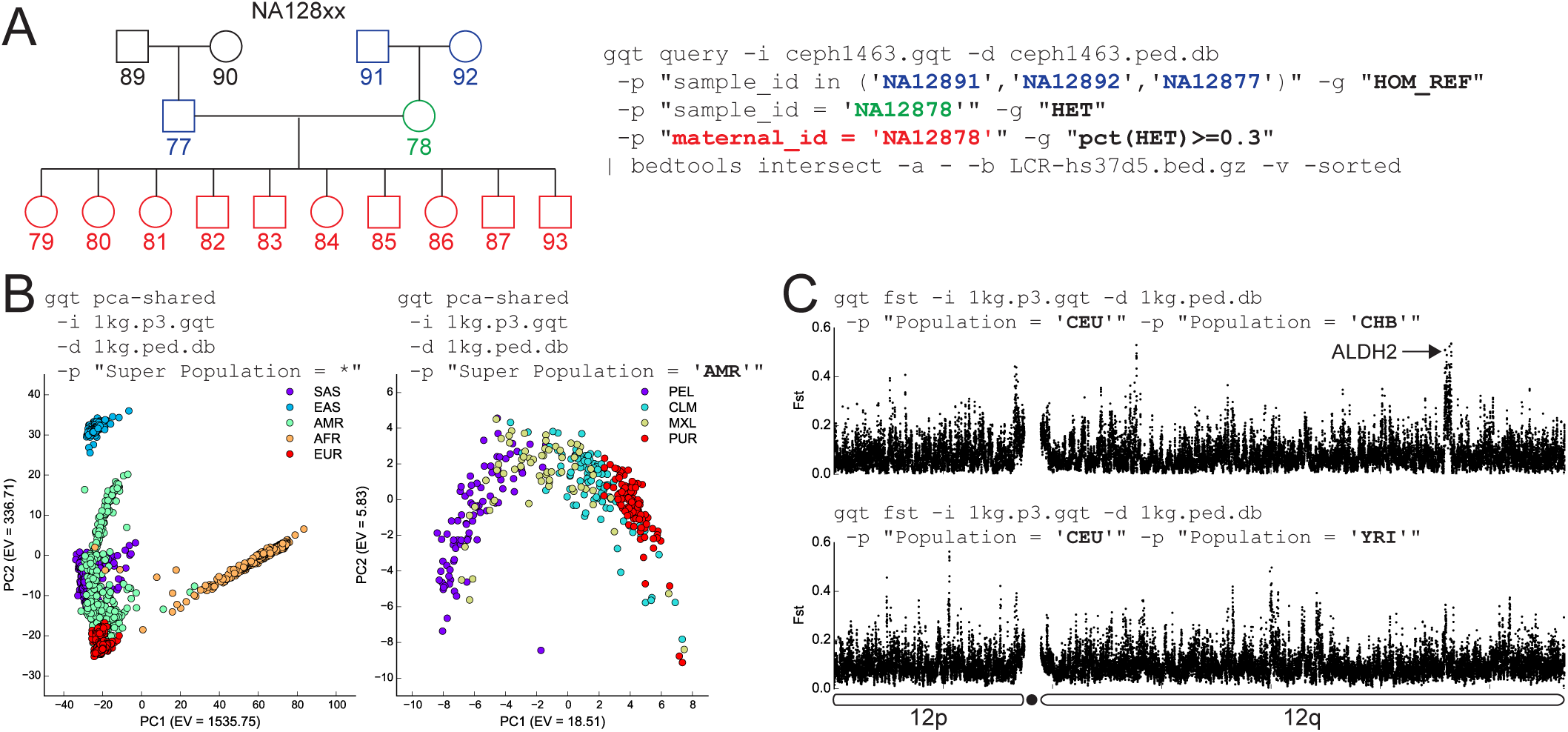
Examples of analysis using the GQT index and query interface. (**A**) A screen for de novo mutations in NA12878 by screening against parental and child genotypes, then filtering those results against low complexity regions using BEDTOOLS. This query reported 2,659 candidates in 0.82 seconds. (**B**) A Principal Components analysis of the variants from Phase 3 of 1000 Genomes requiring 207 minutes for all 2,504 individuals, and 3 minutes for the 347 AMR individuals. (**C**) Fst analysis of Europeans v. East Asians and Europeans v. Africans, where each full genome Fst run was 146X faster than VCFTOOLS. The strongest peak in the East Asian comparison, which was absent in the African comparison, contained the aldehyde dehydrogenase 2 (ALHD2) gene.

#### Fixation index

The fixation index (Fst) is a widely-used metric for summarizing the degree of population differentiation that exists between populations owing to genetic drift or selection. Fst is calculated for each polymorphic locus and the average of per-locus measures across the genome are typically used to summarize the overall genetic structure that exists between populations. Since the Weir and Cockerham^9^ measure is based on the variance in allele frequencies observed between subpopulations, we can exploit the GQT genotype index to compute Fst for all 84.7 million variants in the 1000 Genomes dataset in 74 seconds. This reflects a 146X speedup over the 184 minutes that VCFTOOLS requires to compute Fst for each variant in the Phase 3 dataset. GQT emits individual Fst measures for each polymorphism, and when visualized on an entire chromosome, the patterns of individual measures can reveal loci that are either under selection or genes such as ALDH2, whose haplotypes have markedly different frequencies in different world populations^10^ (Figure 5C).

#### Burden tests

Although the GQT index is structured based on individuals rather than variant loci, it is nonetheless capable of efficiently computing the fundamental measures that underlie many locus-centric statistics such as burden tests. For example, the C-alpha burden test^11^ measures the difference in the number of times an allele is observed in cases and controls versus a binomial expectation of the allele counts in the population as a whole. We have implemented the C-alpha statistic in GQT; once case and control populations are defined via the GQT query interface, a standard summary statistic (in this case minor allele frequency) is computed for both subpopulations at each variant site. The statistics for each variant are then aggregated across a given locus and normalized by the variance to produce the final statistic for each gene or locus. We emphasize that C-alpha is merely a representative example of the several burden tests that follow this computational pattern and are therefore candidates for future integration into the GQT toolset.

## DISCUSSION

Recognizing the scaling challenges posed by current and future genome datasets, our motivation was to explore new data compression and indexing strategies that minimize storage requirements while also enabling highly efficient analyses of the underlying data. We have shown that variant searches driven by sample genotypes and phenotypes are substantially faster when genotype data is transposed from a variant-centric to individual-centric organization. Competitive data compression is achieved by arranging variants in order of allele frequency, followed by word-aligned hybrid encoding of genotype bitmaps. Moreover, because WAH compression of the genotype index minimizes data inflation, GQT also achieves substantially better analysis performance for queries that interrogate the genotypes of the individuals in the dataset.

We have shown how individual-centric data organization can expedite a wide range of statistical and population genetics measures, since many of these statistics fundamentally depend on the comparison of two or more individuals’ genotypes. However, GQT is poorly suited for haplotype-based analytical approaches^12^ and statistical measures^13,14^ because GQT orders variant sites by allele frequency instead of chromosomal position. Nonetheless, WAH-compressed bitmaps are not inherently constrained to this organization. We plan to explore variant-centric bitmap indices to enable efficient screens for selective sweeps and to quickly search for haplotypes that are identical by descent.

While the GQT approach is well suited to very large datasets where many variants are extremely rare (and thus highly compressible), it provides less benefit for datasets involving few variants, samples, or both. This is illustrated in our comparisons to PLINK and BCFTOOLS for smaller subsets of the 1000 Genomes project dataset (Figure 2), and in our analyses (Supplementary Table 1) of variant datasets from the Mouse Genome Project (28 samples, 68.1 million variants) and the Drosophila Genome Reference Panel (205 samples, 6.1 million variants). It is nonetheless important to emphasize that despite this manuscript’s focus on human genomes, the strategies we have devised are generally applicable to large-scale datasets from any organism.

GQT’s current indexing strategy is restricted to queries in search of variants based on unphased sample genotypes, phenotypes, and relationships. We recognize that more general queries will require indices of other genotype metadata such as allele-specific sequencings depths, genotype phase, individual haplotypes, and genotype likelihoods in order to enable a broader range of analyses and to impose stricter quality control over the variants that are returned. Bitmap indices are poorly suited to attributes having continuous values (e.g., genotype likelihoods), since one bit array must be created for each distinct value. However, based on the success of previous studies^15^, binning continuous values into large sets of discrete values is likely to be a straightforward approach to indexing continuous values with minimal information loss. Indeed, such binning is already employed in the VCF specification for Phredscaled^16^ integer representations of genotype likelihoods. We expect the inherent analytical flexibility of the GQT index to be of broad use and we will therefore strive to provide a convenient software interface for its integration into existing toolsets and future method development.

Lastly, since GQT optimizes individual-centric and genotype-focused analyses, it is a natural complement to the variant-centric indexing strategies employed by methods such as Tabix^17^ and BCFTOOLS. Integration of these naturally complementary indexing strategies will provide the basis for a rich query interface supporting genetic data exchange efforts such as the Global Alliance for Genomics and Health. Based on the efficiency and inherent flexibility of this genotype indexing strategy, we expect GQT to be a broadly useful general indexing strategy enabling the exploration of massive genomics datasets involving thousands or even millions of genomes.

## URLS

All source code for the GQT toolkit is available at http://exac.broadinstitute.org/about. Furthermore, all commands used for the experiments conducted in this study are available at https://github.com/ryanlayer/gqt_paper.

## ONLINE METHODS

We compared GQT v0.0.1 to PLINK v1.90p and BCFTOOLS (https://github.com/samtools/bcftools) 1.1 in terms of index file size and query runtime against four large-scale cohorts and simulated data sets. The cohorts included: 2,504 human genomes from the 1000 Genomes Project phase 3, 28 mouse genomes from the Mouse Genomes Project, 205 fly genomes from the *Drosophila* Genetic Reference Panel (DGRP), and 60,706 human exomes from Exome Aggregation Consortium (ExAC). Query comparisons included time to compute the alternate allele frequency count for a target 10% of the population, and time to find rare (details below) variants among a target 10% of the population. Both target sets were comprised of the last 10% of individuals. For all runtime comparisons BCFTOOLS considered a BCF file, PLINK considered a BED and BIM file, and GQT considered a GQT index and BIM file (details in Supplementary Materials). Runtimes for GQT considered two different modes: the default mode that reports all matching variants in full VCF format and the “`count`” mode (specified by the “`-c`” option) that only reports the number of matching variants. The “`count`” mode is a useful operation in practice, and also demonstrates speed without I/O overhead.

### File size

File size comparisons used an uncompressed VCF as a baseline, BCFTOOLS used a compressed binary VCF (BCF) to store both variant and sample data, PLINK used the binary plink format (BED) to store sample data and a BIM file to store variant data, and GQT used a GQT index file to store WAH-encoded sample genotype data as well as a BIM file to store LZ77-compressed variant data.

### Alternate allele count

The baseline runtime for finding the alternate allele count was the BCFTOOLS “`stats`” command with the “`-S`” option to select the subset of individuals, the PLINK command was “`--freq`” with the “`--keep`” option to select individuals, and the GQT command was “`query`” (with and without the “`-c`” option) with the “`-g "count(HET HOM_ALT)"`” option to specify the allele count function and the “`-p "Ind_ID >= N"`” option to select the subset (where N was the ID of the range that was considered).

### Identifying rare variants

The baseline runtime for selecting the variants was the BCFTOOLS “`view`” command with the “`-S`” option to select the subset of individuals and the “`-C`” option to limit the frequency of the variant, and the GQT command was “`query`” (with and without the “`-c`” option) with the “`-g "count(HET HOM_ALT) < =F"`” option to specify the allele count filter (where F was the maximum occurrence of the variant) and the “`-p "Ind_ID > = N"`” option to select the subset (where N was the ID of the range that was considered). In both cases the limit was set to either 1% of the subset size or 1, whichever was greater. PLINK was omitted from this comparison because third-party tools are required to complete this operation, and in our opinion it is not fair assign the runtime of those tools to PLINK.

### Pedigree analyses

GQT can report the variants matching any number of genotype and phenotype queries pairs. Each query pair selects the variants that meet the specified conditions. For example, “`-p "sample_id = 'NA12878'" -g "HET"`” will select only those variants where NA12878 is heterozygous. Any subsequent genotype/phenotype query pair will further restrict the final set of variants to the subset meeting the extra conditions. For example, the query “`-p "sample_id = 'NA12878'" -g "HET" - p "maternal_id = 'NA12878'" -g "pct(HET HOMO_ALT) > =0.3"`” includes two pairs and will return only those variants where NA12878 is heterozygous and where at least 30% of her children are also non-reference.

This analysis considered the full CEPH 1473 pedigree. Since GQT reports results in valid VCF format, we were able to use other existing software to filter the results. In particular, we used BEDTOOLS to filter out any variants fell within a low complexity region as identified by Li^8^.

The resulting GQT command was:

~~~
gqt query -i cohort-illumina-wgs.vcf.gz.gqt -d ceph1463.ped.db \
~~~

- ~~~
-p "sample_id in ('NA12891','NA12892','NA12877')" \
~~~
- ~~~
-g "HOMO_REF" \
~~~
- ~~~
-p "sample_id = 'NA12878'" \
~~~
- ~~~
-g "HET" \
~~~
- ~~~
-p "maternal_id = 'NA12878'" \
~~~
- ~~~
-g "pct(HET HOMO_ALT) > =0.3" \
~~~

~~~
| bedtools intersect -v \
~~~

- ~~~
-a stdin \
~~~

- ~~~
-b btu356_LCR-hs37d5.bed/btu356_LCR-hs37d5.bed \
~~~

~~~
> NA12878_de_novo.vcf
~~~

### Principal component analysis (PCA)

Using the “`pca-shared`” command, GQT computed a score for each pair of individuals in the target population that reflected the number of shared non-reference loci between the pair. This score was calculated in two stages. First, an intermediate OR operation of the `HET` and `HOM_ALT` bitmaps within each individual produced two bitmaps (one for each member of the pair) that marked non-reference loci. Then, an AND of these two bitmaps produced a final bitmap that marked the sites where both individuals were non-reference. GQT then counted the number of bits that were set in this bitmap and reported the final score. The “`pca-shared`” command also takes the target population as a parameter. The first plot in Figure 5B considered all 2504 individuals, which included the South Asian (SAS), East Asian (EAS), Admixed American (AMR), African (AFR), and European (EUR) “ super populations”. The second plot in Figure 5B only considered the 347 individuals in the Admixed American (AMR) super population, which included: Peruvians from Lima, Peru (PEL), Columbians from Medellin, Columbia (CLM), Mexican Ancestry from Los Angeles, USA (MXL), and Puerto Ricans from Puerto Rico (PUR).

This analysis considered only the autosomes within the 1000 Genomes phase 3 variants. The result of the GQT “`pca-shared`” command is the upper half of a square symmetrical matrix. The Python script “`fill_m.py`” fills out a full matrix by reflecting those values, and another Python script “`pca_light.py`” calculates the Eigen vectors and values using the Numpy scientific computing library and plots the results.

The resulting GQT commands were:

~~~
gqt pca-shared \
~~~

- ~~~
-i ALL.phase3.autosome.vcf.gz.gqt \
~~~
- ~~~
-d integrated_call_samples.20130502.ALL.spop.ped.db \
~~~
- ~~~
-p "*" \
~~~
- ~~~
-f "Population" \
~~~
- ~~~
-l ALL.phase3.autosome.vcf.gz.gqt.pops \
~~~

~~~
> ALL.phase3.autosome.vcf.gz.gqt.o
~~~

~~~
gqt pca-shared \
~~~

- ~~~
-i ALL.phase3.autosome.vcf.gz.gqt \
~~~
- ~~~
-d integrated_call_samples.20130502.ALL.spop.ped.db \
~~~
- ~~~
-p "Super_Population = 'AMR'" \
~~~
- ~~~
-f "Population" \
~~~
- ~~~
-l ALL.phase3.autosome.vcf.gz.gqt.AMR.pops \
~~~

~~~
> ALL.phase3.autosome.vcf.gz.gqt.AMR.o
~~~

### Fixation index

Fst is a widely used measurement of the genetic difference between populations, and here we focused on the method proposed by Weir and Cockerham^9^. While this metric has many parameters, it is fundamentally based on the frequency of an allele and the proportion individuals that are heterozygous for that allele in each population. GQT can quickly calculate both metrics for various populations across the whole genome. Allele frequency is calculated by considering the bits marked in both the `HET` and `HOM_ALT` bitmaps. Since each bit in a bitmap corresponds to a specific variant, calculating the allele frequency of all variants involves incrementing the associated counter for each set bit. WAH encoding makes this process more efficient by collapsing large stretches of reference alleles (which are represented by zeros in the `HET` and `HOM_ALT` bitmaps) into a small number of words that can be quickly skipped. We further accelerate the processing of each word by using the AVX2 vector-processing instruction set. Vector processing instructions are set of special CPU instructions and registers that exploit data-level parallelism by operating on a vector of values with a single operation. These instructions allow us to consider 8 bits in parallel, and therefore compute the resulting sum of each word in 4 (as apposed to 32) operations. The proportion of individuals that are heterozygous for an allele is computed in a similar manner, except only the `HET` bitmaps are considered.

This analysis considered the 1000 Genomes phase 3 variants that were bi-allelic and had a minimum alternate allele frequency of 1%. The GQT “`fst`” command can consider two or more populations with the “`-p`” option. The first plot in Figure 5C considered the Utah Residents with Northern and Western European Ancestry (CEU) and the Han Chinese in Beijing, China populations (“`-p "Population = 'CHB'" -p "Population = 'CEU'"`”). The second plot in Figure 5C considered the Utah Residents with Northern and Western European Ancestry (CEU) and the Yoruba in Ibadan, Nigeria populations (“`-p "Population = 'CHB'" -p "Population = 'YRI'"`”). Both plots in Figure 5C were smoothed using the mean Fst value over a 10kb window with a 5kb step. The comparison to VCFTOOLS considered version 0.1.12 and the “`--weir-fst-pop`” options for the CHB and CEU populations.

The resulting GQT commands were:

~~~
gqt fst \
~~~

~~~
-i \
~~~

~~~
ALL.phase3_shapeit2_mvncall_integrated_v5a.20130502.genotypes.vcf.gz.gqt \
~~~

~~~
-d 1kg.phase3.ped.db \
~~~

~~~
-p "Population = 'CHB'" \
~~~

~~~
-p "Population = 'CEU'" \
~~~

~~~
> CHB_vs_CEU.gqt.fst.vcf
~~~

~~~
gqt fst \
~~~

~~~
-i \
~~~

~~~
ALL.phase3_shapeit2_mvncall_integrated_v5a.20130502.genotypes.vcf.gz.gqt \
~~~

~~~
-d 1kg.phase3.ped.db \
~~~

~~~
-p "Population = 'YRI'" \
~~~

~~~
-p "Population = 'CEU'" \
~~~

~~~
> YRI_vs_CEU.gqt.fst.vcf
~~~

### Burden tests

GQT can support gene or locus based burden tests by computing the test parameters for every variant, and reporting the results in position-sorted order (assuming the source VCF file is ordered). With these pre-computed and ordered values, the results can be easily aggregated into groups (e.g., genes) by a second pass of the data. Here, we considered the C-alpha test proposed by Neal in 2011^11^. This test is based on the number of times a variant is observed in cases and controls. GQT uses a process similar to that described in the previous Fixation index section to count the occurrence alleles in case and control populations (where are each specified by a phenotype query). Once these parameters are computed for every variant and reported in position-sorted order, the Python script “`calpha.py`” aggregates the values for a specified set of groups in a given BED file, and reports the test statistic and the asymptotic Z score and P-value.

### Experimental Data sets

- **1000 Genomes phase 3**: Individual chromosome VCF files were retrieved from [2] (last accessed December 10, 2014) and combined into a single file using the BCFTOOLS “`concat`” command. To understand how each tool scaled as the number of samples and variants increased, we subsampled the full data set (which included 2,504 individuals) to create new sets with 100, 500, and 1,000 individuals. To create each data set size, we randomly selected the target number of samples, then used the BCFTOOLS “`view`” command with the “`-s`” option to return just the genotypes of the target samples. We then recomputed the allele frequency of each variant with the BCFTOOLS “`fill-AN-AC`” plugin, and filtered all non variable sites with the BCFTOOLS “`view`” command and the “`-c 1`” option.
- **Mouse Genomes Project**. Data was retrieved in VCF format from (ftp://ftp-mouse.sanger.ac.uk/current_snps/mgp.v4.snps.dbSNP.vcf.gz) and was last accessed November 25, 2014.
- **Drosophila Genetic Reference Panel**. Data was retrieved in VCF format from (http://dgrp2.gnets.ncsu.edu/data/website/dgrp2.vcf) and was last accessed November 25, 2014.
- ***Exome Aggregation Consortium* (ExAC)**. Version 3 of the ExAC dataset was analyzed and run times were measure on the computing infrastructure at the Broad Institute.
- ***CEPH 1473 pedigree***. A VCF file of variants in the CEPH 1473 pedigree that was sequenced as part of the Illumina Platinum Genomes Project was downloaded from ftp://ftp-trace.ncbi.nih.gov/giab/ftp/data/NA12878/variant_calls/RTG/cohort-illumina-wgs.vcf.gz.

### Simulated data sets

Genotypes were simulated using the MaCS^18^ simulator version 0.5d with the mutation rate and recombination rate per site per 4N generations set to 0.001, and the region size set to 100 megabases. Since our simulation considered between 100 and 100,000 diploid samples, and MaCS only simulates haplotypes, we simulated 2X haplotypes for each case and combined 2 haplotypes to create a single diploid genome. It was computationally prohibitive to produce a data set for 1 million individuals (the 100,000 simulation ran for over four weeks), so we used a simple linear fit to estimate the file size and runtimes for 1 million individuals.

### Computing environment

GQT is a tool written in C, which uses the htslib (https://github.com/samtools/htslib) to interact with VCF and BCF files and zlib (http://www.zlib.net/) to compress and inflate variant metadata. All experiments were run on Ubuntu Linux v3.13.0-43, with gcc v4.9.2, 4 Intel Core i7-4790K 4.00GHz CPUs with the Haswell microarchitecture, and a 550 MB/s read/write solid-state hard drive.

## ACKNOWLEDGEMENTS

We are grateful to Colby Chiang for conceptual discussions, Igor Levicki for helpful advice on AVX2 operations, both Shane McCarthy and Petr Danecek for their guidance with `htslib`, and Zev Kronenberg for advice on population genetics measures. The authors would also like to thank the Exome Aggregation Consortium and the groups that provided exome variant data for comparison. A full list of contributing groups can be found at http://exac.broadinstitute.org/about. This research was supported by an NHGRI award to A.R.Q. (NIH R01HG006693).

## SUPPLEMENTARY METHODS

### Sorting by allele frequency to improve compression

Long runs of identical genotypes are easily compressed. We have chosen an alternative individual-centric data organization strategy that, while it facilitates queries based on individual genotypes, destroys the inherent compression of the genotype runs in the variant-centric approach. This loss of compression is the direct consequence of the fact that records in the individual-centric approach reflect the genotypes for a given individual at each site of variation in the genome. Runs of identical genotypes are far shorter, on average, than with the variant-centric approach and therefore, the individual-centric strategy will yield poor compression. The question then becomes how to leverage the query efficiency of individual-centric data organization while also retaining the opportunity for data compression? GQT solves this by sorting the variant columns of the transposed matrix in order of allele frequency. This results in fewer, longer runs of identical genotypes in each individual’s row of genotypes (**Supplementary Figure 1**). For example, we compared the effect of sorting variants on genotype runs using chromosome 20 from Phase 3 of the 1000 Genomes Project. As expected, sorting variants by allele frequency caused both a dramatic increase in the mean length (10.7 versus 23.2) of identical genotype runs and a reduction in the median number of runs per individual (158,993.5 versus 70,718.5). Fewer, longer identical genotype runs allows greater compression of the each individual’s (reordered) variant genotypes.

**Supplementary Figure 1.**
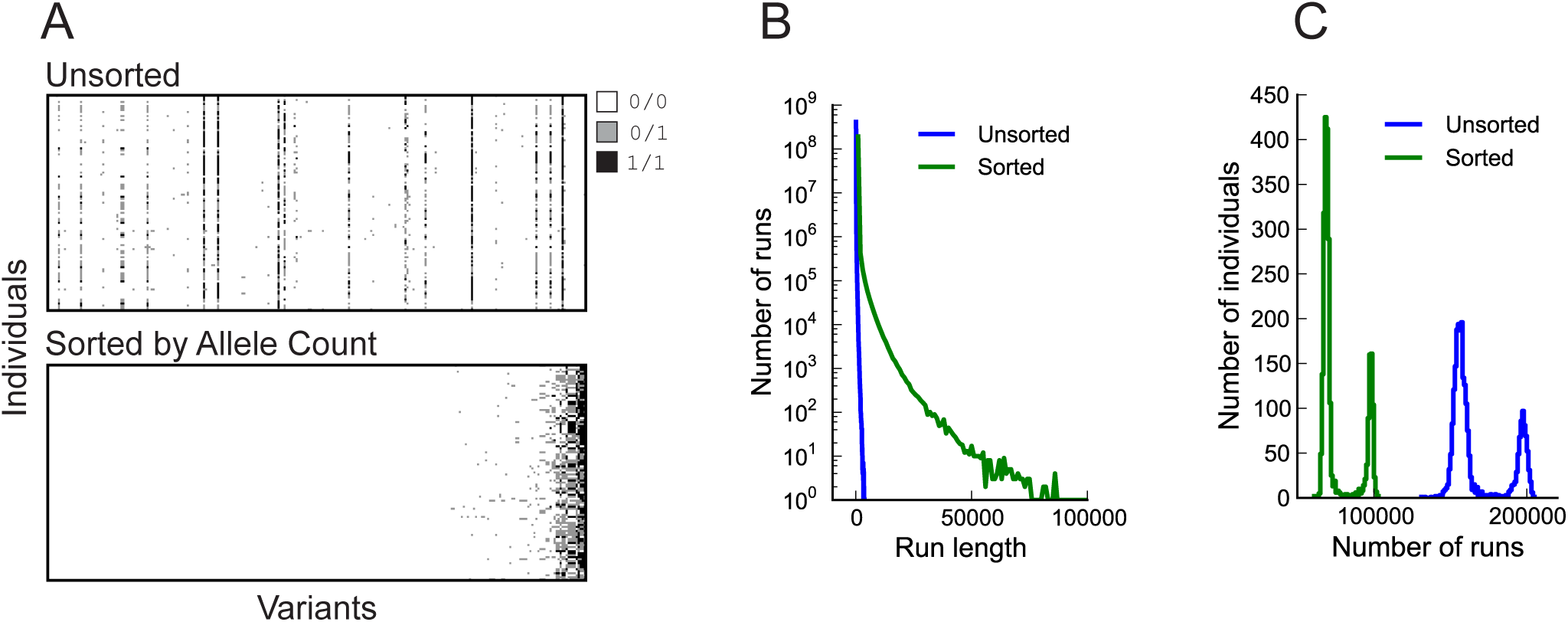
Sorting variants by allele frequency improves compression. (**A**) A graphical comparison of the genotype distribution of individuals (rows) and variants (columns) before and after sorting. These data represent genotypes from the 1000 Genomes Project, phase 3, for a portion chromosome 20. (**B**) The run length distribution for unsorted and sorted genotypes. (**C**) The distribution of the number of runs for sorted and unsorted data. In both cases the second peak is composed predominantly of individuals from African decent (AFR); 604/661 AFR are in the second peak in the sorted case, and 640/661 AFR in the unsorted case.

### Representing sample genotypes with bitmap indices

The fundamental advantage of individual-centric data organization is the fact that all of an individual’s genotypes can be accessed at once. This enables algorithms to quickly compare all variant genotypes from multiple samples in search of variants that meet specific inheritance patterns, allele frequencies, or enrichment among subsets of individuals. Despite the improved data alignment, comparing sample genotypes can still require substantial computation. For VCF, which encodes diploid genotypes as “`0/0`” for homozygotes of the reference allele, “`0/1`” for heterozygotes, “`1/1`” for homozygotes of the alternate allele, and “`./.`” for unknown genotypes (**Supplementary Figure 2A**), comparing the genotypes of two or more individuals requires iterative tests of each genotype for each individual.

**Supplementary Figure 2.**
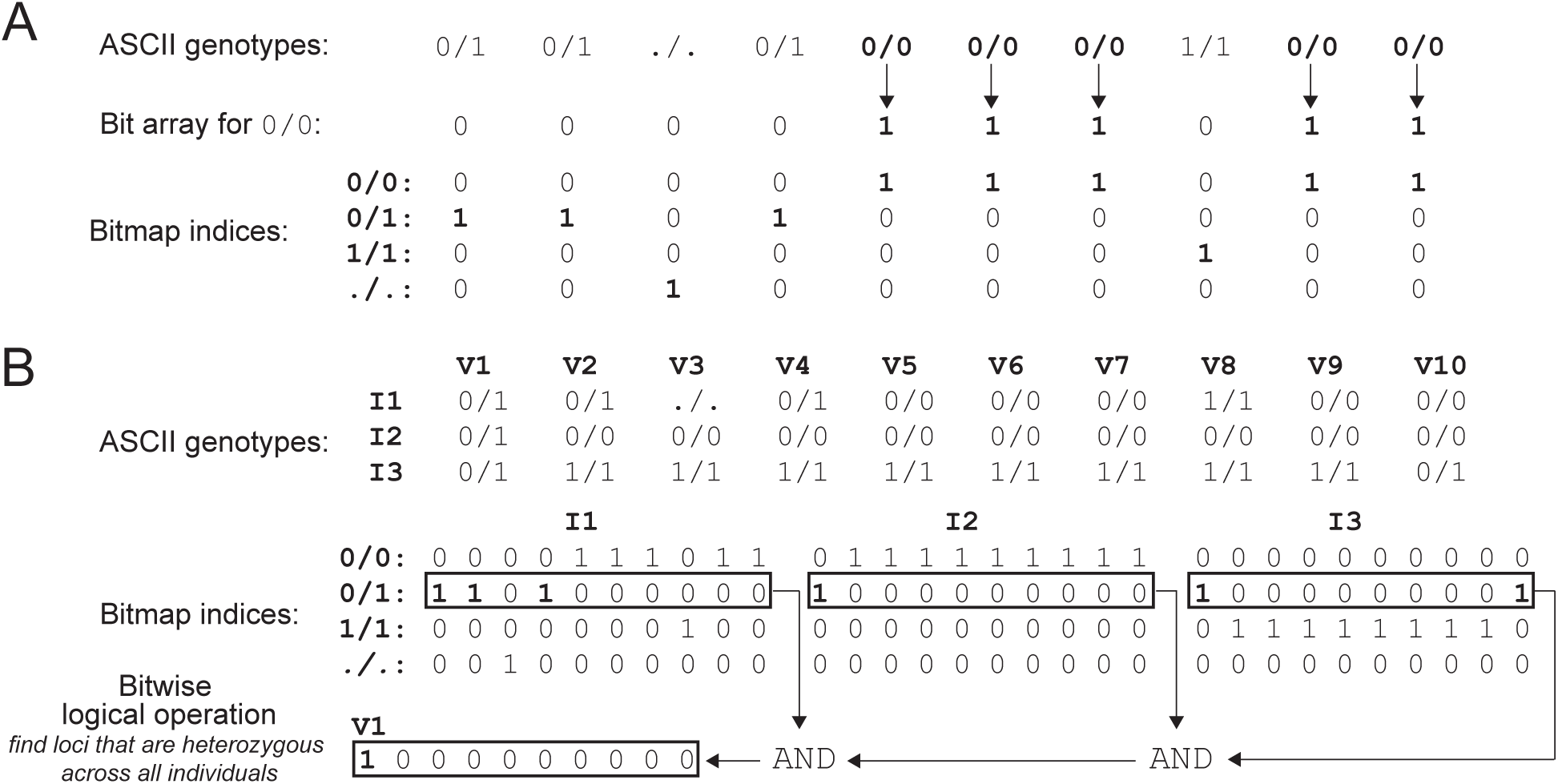
Bitmaps enable rapid genotype comparisons en masse. (**A**) A bit array marks the existence of one genotype state (for example, homozygous reference; “`0/0`”) for all variants. Similarly, a bitmap index is composed of a distinct bit array for each possible genotype state. (**B**) Example genotypes in VCF format are presented for three individuals (I1, I2, I3) at 10 variant sites (V1-V10). A bitwise AND of the bit arrays corresponding to the heterozygous genotype yields the variant that is heterozygous in all individuals.

Recognizing this inefficiency, we encode each individual’s vector of genotypes with a bitmap index. A Bitmap index (“bitmap”) is an efficient strategy for indexing attributes with discrete values that uses a separate bit array for each possible attribute value. In this case of an individual’s genotypes, a bitmap is comprised of four distinct bit arrays corresponding to each of the four possible (including “unknown”) diploid genotypes. As illustrated in **Supplementary Figure 2A**, the bits in each bit array are set to true (`1`) if the individual’s genotype at a given variant matches the genotype the array encodes. Otherwise, the element is set to false (`0`). In turn, bitmap encoding facilitates the rapid comparison of individuals’ genotypes with highly optimized bitwise logical operations. As an example, a bitmap search for variants where all individuals are heterozygous involves a series of pairwise `AND` operations on the entire heterozygote bit array from each individual. The intermediate result of each pairwise `AND` operation is subsequently compared with the next individual, until the final bit array reflects the variants where all individuals are heterozygous (**Supplementary Figure 2B**). Such queries are expedited owing to the ability of CPUs to simultaneously test multiple bits (i.e., genotypes) with one bitwise logical operation.

### Efficient comparisons using bitmaps

By using bitwise logical operations we can compare many genotypes in a single operation, rather than comparing each individual value. At a low level, bitwise logical operations are performed on words, which are the fixed-size unit of bits used by the CPU. Modern processors typically use either 32- or 64-bit words. When a bitwise logical operation is performed between two bit arrays (each of which correspond to the genotypes of two individuals), the processor completes this operation on one pair of words at a time. For example, if the word size is 32, then computing the bitwise `AND` of two bit arrays that are each 320 bits long would require only 10 `AND`s. This optimization is equivalent to a 32-way parallel operation with zero overhead. This concept is demonstrated in **Supplementary Figure 3**, where we are searching for the loci where all three individuals are heterozygous among eight variants. When genotypes are encoded in ASCII (as they are in VCF), the algorithm must loop over every individual and every variant to find the common sites. In total, this requires 24 iterations (**Supplementary Figure 3A**). In contrast, encoding genotypes with a bitmap allows the same computation to be completed with only three bit-wise `AND` operations (**Supplementary Figure 3B**). In effect, bit-wise logical operations compare all eight genotypes in parallel in a single step. For brevity an 8-bit word is used, and only the heterozygous bit arrays are show, but the same principles hold for the larger word sizes employed by GQT.

**Supplementary Figure 3.**
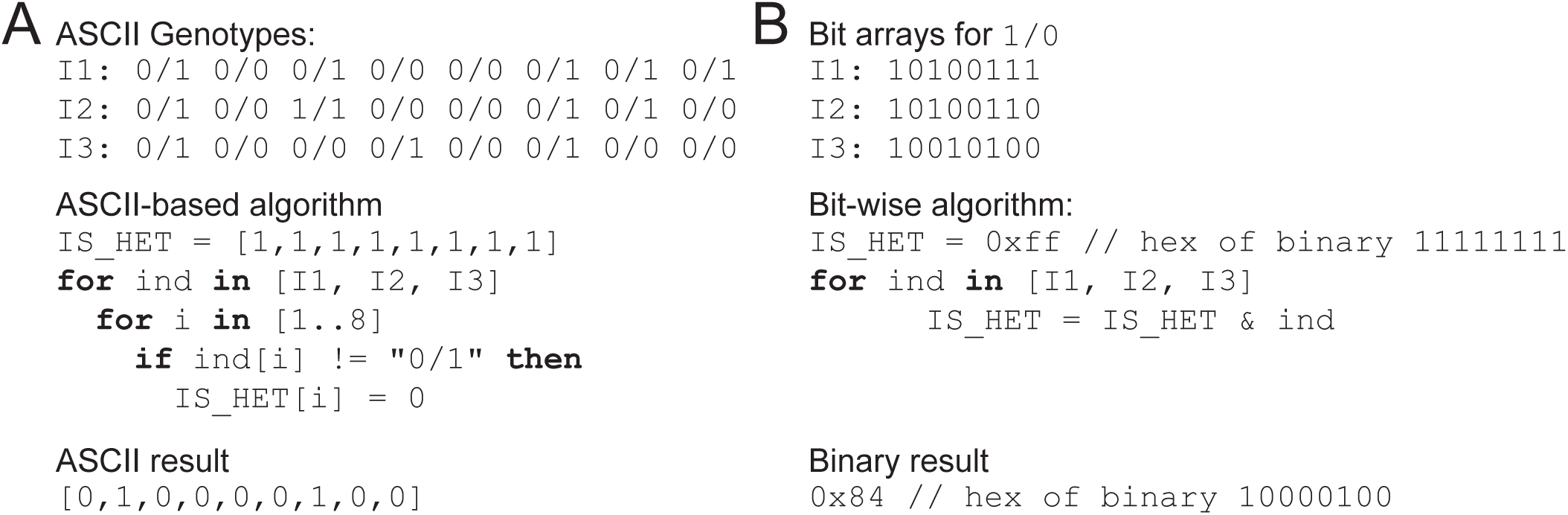
Efficiency improvements for genotype comparisons when using a bitmap index. (**A**) When considering genotypes in ASCII format (e.g., VCF), an algorithm searching for the set of variants that are heterozygous in all individuals must operate on every genotype for each for every individual separately. (**B**) In contrast, when genotypes are represented with a bitmap index, where a set of genotypes are encoded into a single CPU word (for brevity, only the bit arrays associated with the heterozygous state are shown), bitwise logical operations can be used to operate on all of the genotypes in the word with a single operation. This example assumes a word size of 8, but modern CPU support up to a 64-bit word. For the 24 genotypes given here (3 individuals, 8 genotypes each), the ASCII-base algorithm executes the “`if`” statement 24 times, while the bit-wise algorithm executes the logical AND (“`&`”) only three times, with both algorithms producing equivalent results.

### Using Word-Aligned Hybrid to directly query compressed data

While bitwise logical operations can drastically improve query runtime performance, bitmap indices require double the amount of space over the minimum two bits per genotype. To address this issue we can look to compressing that data. While genotype data can be compressed with standard run-length encoding, bitwise logical operations require that the bits associated with variants must be aligned, which is difficult to ensure with run-length encoding (RLE). Instead we use the Word Aligned Hybrid (WAH) encoding strategy, which represents run length in words not in bits. As shown in **Supplementary Figure 4A**, RLE encodes stretches of identical values (“ runs”) to a new value where the first bit indicates the run value and the remaining bits give the number of bits in the run. WAH is similar to RLE, except that it uses two different types of values. The “fill” type encodes runs of identical values, and the “ literal” type encodes uncompressed binary. This hybrid approach address an inefficiency in RLE where short runs map to larger encoded values. The first bit in a WAH value indicates whether it is a fill (`1`) or literal (`0`). For a fill, the second bit gives the run value and the remaining bits give the run length in words (not bits, like in RLE). For a literal, the remaining bits directly encode the uncompressed input. Since each WAH-encoded value represents some number of words, and bitwise logical operations are performed between words, these operation can be performed directly on compressed values.

The algorithm that performs bitwise logical operations is straightforward (**Supplementary Figure 4B**). To operate on two uncompressed bit arrays we simply move in unison between the two arrays from the first to last word, and find the result of each pair of words. Since each WAH value encodes one or more words, we need to move across each WAH array independently. At each step we track the number of words that have been considered in the current value and only move to the next word once the words have been exhausted. Bitwise logical operations improve the performance of most queries by computing many genotype comparisons in parallel. Some queries cannot be resolved by these operations alone.

**Supplementary Figure 4.**
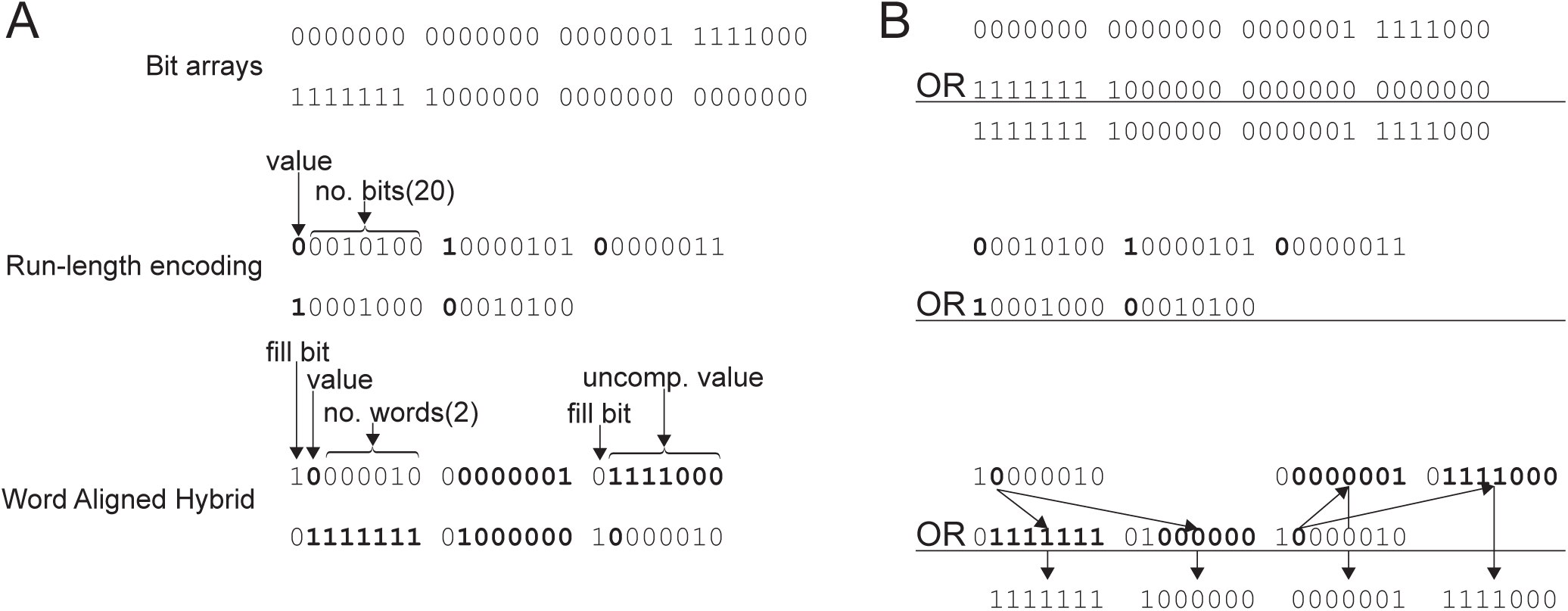
A comparison of binary encodings and associated bit-wise logical operations. (**A**) Two bit arrays are given in three different binary encodings: the uncompressed bit array on top, followed by the run-length encoding (RLE) and word-aligned hybrid encoding (WAH). RLE maps each set of consecutive bits to a new value that uses one bit for the run value, and the remaining bits for the number of bits in the run. WAH maps bits to one of two types of values: those that include runs and those that encode the raw binary. The first bit in a WAH value indicates if the value encodes a run (the fill bit). If that bit is set then the second bit gives the run value and the remaining bits give the run length in number of ***words*** (i.e., not number of ***bits*** as in RLE). If the fill bit is not set then the remaining bits are the uncompressed binary values. (**B**) The logical OR for the three encodings is given. For the uncompressed binary, the OR follows bit for bit across both values. For RLE, the logical OR is undefined (without inflation) because the two encoded values are no longer aligned and have different lengths. For WAH, the values are aligned based on their run length, then the logical OR is performed (in this case) between the value bit and the associated uncompressed values.

Finding the allele frequency among a set of individuals is one such query. In this case, each bit corresponds to the genotype of a particular individual at a particular genomic position, and the allele frequency for that position is the summation of the corresponding bits across all individuals. Since 32 bits are packed into a single word, this process can be reduced to a series of bitwise sums between words, which, unfortunately, is not a standard operation. While no architecture provides single-instruction support for bitwise sum, the operation does exhibit a high-degree of parallelism. The sum of each position is independent of all other positions, which allows (in principle) the sum of all positions to be found concurrently. This classic Single Instruction Multiple Data (SIMD) scenario can be exploited through the use of the vector processor registers and instructions that are supported by the most recent Intel CPUs (Haswell and beyond). These special registers are designed to perform instructions on a list of values in parallel, and by combining several instructions (logical shift, AND, and sum) from the AVX2 instruction set we can perform the bitwise sum of 8 words in parallel. While the 8-way parallelism lags behind what is possible for other operations, it still represents a significant speed up for an operation that is expected to be part of many queries. The index we describe above has the ability to identify variants that meet a complex set of conditions among millions of individuals and billions of genotypes in seconds.

### Contents of BCF file, PLINK index, and GQT index

Direct compression comparisons must account for the fact that each tool compresses different subsets of the variant and sample genotype sections of a VCF file (**Supplementary Figure 5**). By default, the BCF format encodes all of the data and metadata in both sections into binary values, and then compresses those values using blocked LZ77 encoding. PLINK ignores both variant and sample genotype metadata, does not compress the variant data, and simply encodes each genotype with two bits without compression. GQT uses a hybrid strategy for compressing VCF files. It retains all of the variant data and only the genotype values (no metadata) in the genotype section, and stores that data in a “ BIM” file. The variant data is compressed with LZ77 encoding. In addition, an index of individual genotypes is created by transposition, sorting and WAH-compression. Since the variants in the BIM file are stored in the same order as the original VCF, and the genotype index columns are ordered by allele frequency, we must maintain a mapping between these two orderings to retain the ability to print results in the same order as the original VCF. This mapping is stored in a “ VID” file.

**Supplementary Figure 5.**
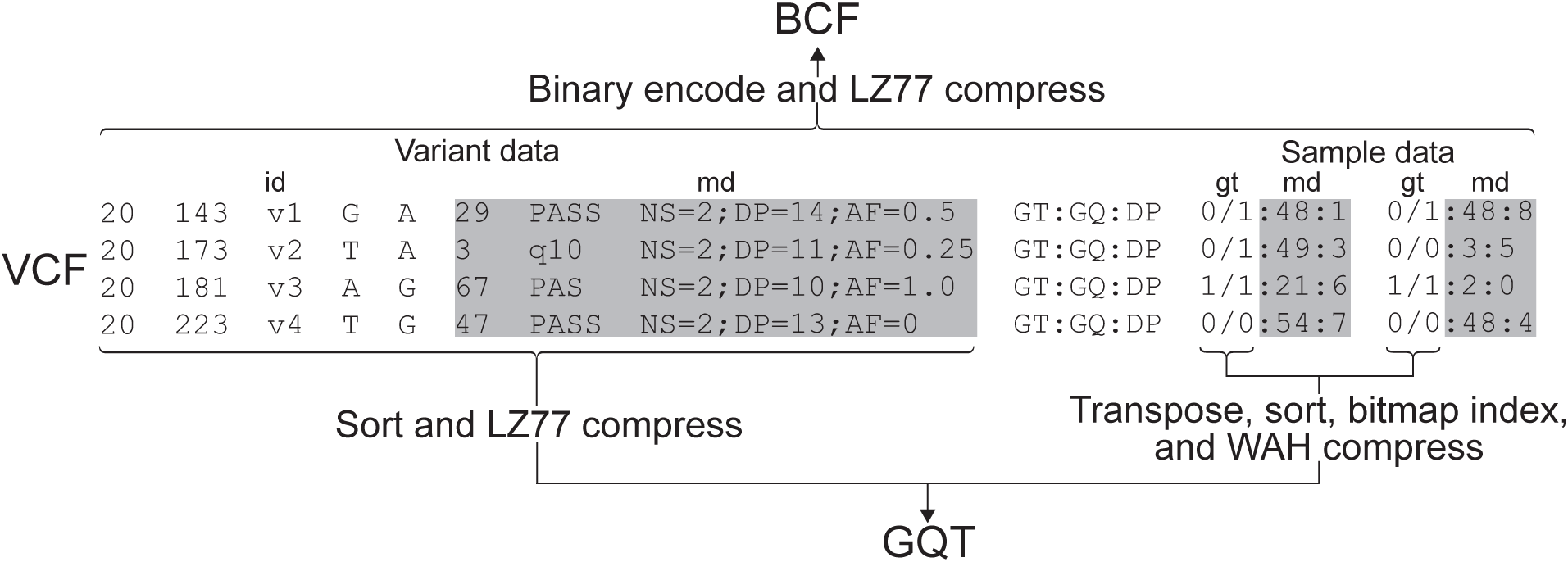
BCF and GQT file composition. VCF files are composed of a variant data section and a sample genotype section and each of these include both core information (e.g. variant position and alleles, and genotype, respectively) and extra metadata. BCF encodes both sections into a binary format, and then compresses the data using blocked LZ77 compression. GQT uses three files: a BIM file that contains all variant data compressed with LZ77, a GQT file with WAH-compressed genotypes (without metadata), and a VID file that stores the mapping between the allele-frequency variant order and the original source VCF variant order. PLINK (not shown) omits all metadata, storing uncompressed binary encoded for genotypes in a BED file and variant positions in a BIM file.

### 32-bit WAH word size

A fundamental choice for WAH-encoding bit arrays is the word size. Modern processors support up to 64 bits, but smaller words of 32, 16, and 8 are also possible, and the choice will affect both the compression ratio and query runtime. Since WAH uses one bit of each word to indicate the type of word (fill or literal), it would seem that larger words would be more efficient. An eight-bit word will have seven useful bits to every overhead bit, while a 64-bit word will have a 63:1 ratio. However, there is a significant amount of waste within fill words. Considering that the first two bits of a fill indicate the word type and run value, and the remaining give the length of the run in words, a 64-bit fill word can encode a run that is 1.4e20 bits long. That is enough bits to encode 46.1 billion human genomes. In fact, we only need 27 bits to cover the full genome, meaning that every 64-bit fill word would have at least 35 wasted bits. This would seem to indicate that smaller words are more efficient, but as the word size decrease the speed up of the bit-wise logical operations also declines. A single operation between two 64-bit words compares 32x more bits (and their associated genotypes) than an operation between two 8-bit words. Taken together, our test show that 32-bits gives the best balance between size and speed.

**Supplementary Figure 6.**
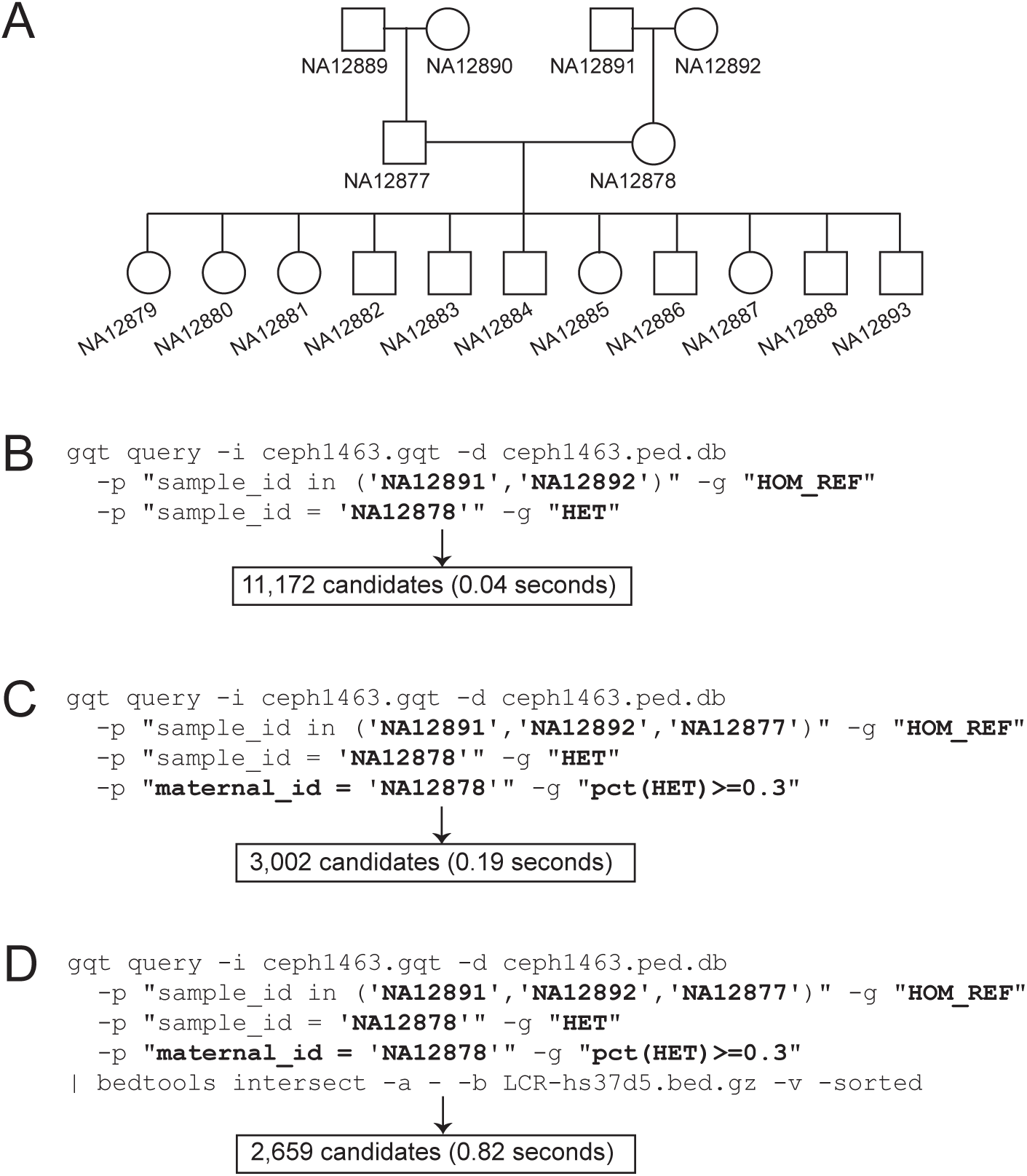
De novo mutation discovery in the CEPH 1463 pedigree. (**A**) The CEPH 1463 pedigree. Our analysis is focused on the discovery of de novo mutations in NA12878, the daughter of NA12891 and NA12892. (**B**) A GQT query for de novo mutations based on the expected genotypes (homozygous for the reference allele) in NA12878’s parents, as well as an expected heterozygous genotype in NA12878. (**C)** True de novo mutations in NA12878’s germline should be passed on to 50% of her offspring, on average. Allowing for genotyping error and binomial expectation, we filter for more confident de novo mutation candidates by requiring that the apparent mutation allele is passed on to at least 30% of NA12878’s children. (**D**) A GQT query that further filters candidate mutations by excluding those lying in low complexity regions of the genome.

**Supplementary Table 1.** The performance of BCFTOOLS, PLINK, and GQT across four cohorts of different sizes and sample populations. This analysis included three whole-genome data sets from three different species: mouse genome project (MGP), *Drosophila* genome reference panel (DGRP), human from 1000 Genomes phase 3 (1KGp3), and a whole-exome for human from the Exome Aggregation Consortium (ExAC).

| Experiment | Tool | MGP<br>(28 samples;<br>68.1 million<br>variants) | DGRP<br>(205 samples;<br>6.1 million<br>variants) | 1KGp3<br>(2,504 samples;<br>84.7 million<br>variants) | ExAC<br>(60,706 samples;<br>9.3 million<br>variants) |
| --- | --- | --- | --- | --- | --- |
| File size (Gb) | BCFTOOLS | 14.94 | 2.99 | 138.41 | 2119.7 |
|  | PLINK | **2.37** | 0.47 | 55.10 | 142.29 |
|  | GQT | 2.97 | **0.43** | **14.33** | **27.96** |
| Alternate allele<br>count time<br>(min.) | BCFTOOLS | 2.79 | 0.43 | 14.30 | 544.30 |
|  | PLINK | **0.41** | 0.05 | 2.61 | 20.10 |
|  | GQT | 1.07 | **0.04** | **0.97** | **1.50** |
| Rare variant<br>search time<br>(min.) | BCFTOOLS | 4.44 | 0.64 | 39.34 | 931.40 |
|  | PLINK | NA | NA | NA | NA |
|  | GQT | **0.75** | **0.03** | **0.86** | **2.10** |

## REFERENCES

1. Zuk, O. et al. Searching for missing heritability: Designing rare variant association studies. Proc. Natl. Acad. Sci. 111, E455–E464 (2014).

2. Purcell, S. et al. PLINK: a tool set for whole-genome association and population-based linkage analyses. Am. J. Hum. Genet. 81, 559–575 (2007).

3. Danecek, P. et al. The variant call format and VCFtools. Bioinforma. Oxf. Engl. 27, 2156–2158 (2011).

4. Keinan, A. & Clark, A. G. Recent Explosive Human Population Growth Has Resulted in an Excess of Rare Genetic Variants. Science 336, 740–743 (2012).

5. 1000 Genomes Project Consortium et al. An integrated map of genetic variation from 1,092 human genomes. Nature 491, 56–65 (2012).

6. Kesheng Wu, Otoo, E. J. & Shoshani, A. Compressing bitmap indexes for faster search operations. in 99–108 (IEEE Comput. Soc, 2002). doi:10.1109/SSDM.2002.1029710

7. Quinlan, A. R. & Hall, I. M. BEDTools: a flexible suite of utilities for comparing genomic features. Bioinformatics 26, 841–842 (2010).

8. Li, H. Toward better understanding of artifacts in variant calling from high-coverage samples. Bioinformatics 30, 2843–2851 (2014).

9. Weir, B. S. & Cockerham, C. C. Estimating F-Statistics for the Analysis of Population Structure. Evolution 38, 1358 (1984).

10. Oota, H. et al. The evolution and population genetics of the ALDH2 locus: random genetic drift, selection, and low levels of recombination. Ann. Hum. Genet. 68, 93–109 (2004).

11. Neale, B. M. et al. Testing for an Unusual Distribution of Rare Variants. PLoS Genet. 7, e1001322 (2011).

12. Sabeti, P. C. et al. Detecting recent positive selection in the human genome from haplotype structure. Nature 419, 832–837 (2002).

13. Voight, B. F., Kudaravalli, S., Wen, X. & Pritchard, J. K. A Map of Recent Positive Selection in the Human Genome. PLoS Biol. 4, e72 (2006).

14. Sabeti, P. C. et al. Genome-wide detection and characterization of positive selection in human populations. Nature 449, 913–918 (2007).

15. Hsi-Yang Fritz, M., Leinonen, R., Cochrane, G. & Birney, E. Efficient storage of high throughput DNA sequencing data using reference-based compression. Genome Res. 21, 734–740 (2011).

16. Ewing, B. & Green, P. Base-calling of automated sequencer traces using phred. II. Error probabilities. Genome Res. 8, 186–194 (1998).

17. Li, H. Tabix: fast retrieval of sequence features from generic TAB-delimited files. Bioinforma. Oxf. Engl. 27, 718–719 (2011).

18. Chen, G. K., Marjoram, P. & Wall, J. D. Fast and flexible simulation of DNA sequence data. Genome Res. 19, 136–142 (2008).

